# Standardizing mechanical dose delivery to cells via nanogroove-guided alignment

**DOI:** 10.64898/2026.08.25.747069

**Authors:** Luigi Crimaldi, Valerio Rosiello, Carlo F. Natale, Valeria Panzetta, Paolo A. Netti

## Abstract

The development of novel mechanomedicine technologies critically depends on the ability to administer a well-defined mechanical dosage to cells. Unlike chemical cues, mechanical signals are vectorial rather than scalar, making their precise delivery inherently complex. When external mechanical stimuli are applied to cells seeded on a flat substrate, the mechanical dose experienced by each cell varies depending on its orientation and conformation, rendering consistent and effective mechano-modulation impractical. Here, we introduce a substrate-guided mechanical stimulation strategy that standardizes mechanical dose delivery at the population level by controlling cell orientation. Using nanogrooved PDMS substrates integrated into a uniaxial stretching platform, we induced coherent alignment of NIH3T3 fibroblasts and their mechanosensitive subcellular structures along the direction of applied strains. Cells cultured on flat or nanogrooved substrates were subjected to sustained uniaxial strains of 8% and 29%, and their responses were quantified in real time by live-cell fluorescence imaging. Nanogroove-induced alignment enabled uniform transmission of substrate strain to focal adhesions and the cytoskeleton, resulting in coherent and quantifiable nuclear deformation across the cell population. In contrast, cells on flat substrates exhibited orientation-dependent deformation modes that canceled out at the population level, leading to heterogeneous and attenuated responses. While cellular adaptation to sustained strain was primarily governed by strain magnitude, substrate-guided alignment markedly reduced cell-to-cell variability in mechanical signal perception. Overall, this work establishes cell alignment as a key parameter for standardizing mechanical dose delivery and improving the reproducibility of mechanobiology experiments and the design of mechanically active biomaterials.

## 1. Introduction

In their native environment, cells are continuously subjected to physical cues that significantly influence their functions in both physiological and pathological contexts. Among these cues, mechanical signals arising from the extracellular matrix (ECM) play a central role in governing key cellular functions such as migration, proliferation and differentiation^1–5^. Cells establish a dynamic mechanical equilibrium with the surrounding ECM, and perturbation of this balance can profoundly alter cell functions. Over the past decades, the field of mechanobiology has sought to elucidate how externally applied mechanical stimuli are sensed by cells and converted into biochemical signals that ultimately regulate gene expression and cell fate^6^. Central to this process are focal adhesions (FAs), multiprotein complexes that physically couple the ECM to the actin cytoskeleton and act as primary mechanosensory sites^7,8^. Mechanical cues transmitted through FAs modulate cytoskeleton organization and intracellular tension, which are subsequently relayed to the nucleus via the linker of nucleoskeleton and cytoskeleton (LINC) complex^4,9^. Nuclear deformations resulting from these forces can alter chromatin organization and transcriptional activity, thereby activating or repressing mechanosensitive genes^10,11^. Importantly, mechanotransduction is not a static process. Cells dynamically adapt to the mechanical perturbation by remodelling FAs, reorganizing the cytoskeleton, and adjusting intracellular tension to maintain mechanical homeostasis^12,13^. Through this adaptive response, external forces can reshape the mechanical identity of cells, which is a key determinant of their functional state and fate^14,15^.

In pharmacology, cell-based drug screening relies on the concept of a well-defined dose (typically expressed as drug concentration relative to cell number) to establish a dose-response relationship^16^. By contrast, in mechanobiology, defining and administering a reproducible “*mechanical dose*” remains a major challenge. Unlike biochemical stimuli, mechanical forces are vectorial quantities characterized by magnitude, direction and plane of application. Consequently, delivering a uniform and controllable mechanical stimulus across a cell population is intrinsically more complex.

To address this challenge, significant efforts have been devoted to the development of in vitro platforms capable of applying mechanical forces to cells, including shear stress in microfluidic systems, cyclic stretching of elastic substrates, and compressive loading devices^17–21^. These approaches have demonstrated the ability to modulate a wide range of cellular behaviour, from endothelial alignment to stem cell differentiation. However, despite these advances, establishing a direct and reproducible relationship between the applied mechanical stimulus and the resulting cellular response remains elusive. A major, yet often overlooked, source of variability arises from the relative orientation between individual cells and the direction of applied force. Conventional culture substrates do not impose any spatial organization on adherent cells, which therefore adopt random orientation. As a result, cells within the same population experience different effective components of the applied mechanical stimulus, leading to highly heterogeneous mechanobiological responses. Indeed, it has been shown that cellular response to substrate stretching is strongly anisotropic: cells aligned parallel or perpendicular to the stretching direction exhibit markedly different FA dynamics, cytoskeleton remodelling and nuclear deformation^22–28^. This intrinsic heterogeneity fundamentally limits the reproducibility of mechanobiology experiments and hampers the establishment of meaningful mechanical dose-response relationship. Without control over cell orientation, population-averaged measurements may obscure or even cancel out biologically relevant responses occurring at the single-cell level. In this context, substrate topography has emerged as an effective tool to control cell orientation through contact guidance^27–29^. Micro- and nanoscale patterns can spatially constrain FAs formation, promote cytoskeleton alignment, and ultimately orient the entire cell body along a predefined direction. By imposing a common reference frame across a cell population, topographical cues offer a promising approach to standardise the transmission and perception of externally applied mechanical forces.

Here, we introduce a cell stimulation strategy that integrates substrate nanotopography with uniaxial mechanical stretching to achieve consistent and population-wide mechanical dosing. Using nanogrooved (Ng) PDMS substrates, we induced uniform alignment of NIH3T3 fibroblasts and their mechanosensitive subcellular structures along the direction of applied strains. Cells cultured on Ng or flat substrates were subjected to sustained uniaxial strains of 8% and 29%, and their response was analysed in real time using live fluorescence imaging. By quantitatively examining FA dynamics, cytoskeleton organization and nuclear deformation as a function of cell orientation, strain magnitude, and time, we demonstrate that nanogroove-induced alignment markedly reduces variability in mechanical signal transmission. This approach enables coherent nuclear deformation across the cell population and reveals strain-dependent adaptive responses that are otherwise obscured on conventional flat substrates. Collectively our findings establish substrate-guided cell alignment as a practical and robust strategy to standardize mechanical dose delivery in mechanobiology experiments.

## 2. Materials and methods

### 2.1 Cell stretching system

The cell stretching chamber was entirely realized in polydimethylsiloxane (PDMS) (Sylgard 184, Dow Corning Corporation, Michigan, USA) through the attachment of a chamber well to the cell culture membrane (Fig. S1A). PDMS base and curing agent were mixed at 10:1 weight ratio and degassed under vacuum for 1 h. Flat and Ng membranes with a thickness of ∼100 µm were produced by spin coating PDMS solution at 500 rpm for 1 min. Nanogroove patterned substrates were obtained by replica molding of PDMS on polycarbonate master^28,30^. The pattern consisted of parallel nanogrooves with a groove and ridge width of 700 nm and depth of 200 nm (Fig. S2A-B). PDMS solution was degassed, poured onto the masters and then cured at 60°C for 2 h. A smooth poly(methyl methacrylate) (PMMA) sheet was implemented to obtain flat membranes curing at 70 °C for 2 h. The chamber well was fabricated by pouring PDMS solution into a 3D printed replica master (3D printer Stratasys Object 30). After curing at 80 °C for 2 h, the PDMS chamber well was subsequently detached from the master and attached to the membrane. The bonding was realized using uncured PDMS as an adhesive glue at the interface between the base of the chamber well and the membrane and curing the assembled cell chamber at 80 °C for 2 h (Fig. S2C).

The realized PDMS cell chamber (49 × 49 × 7 mm^3^) has a central bottom, with dimensions of 32 × 15 × 0.1 mm^3^, on which cells are cultured (Fig. S1B). The uniaxial cell stretching is performed symmetrically pulling the chamber by two ends. Each of the latter is composed of three holes. The two lateral holes are used to perform the traction. The central hole, instead, is used to integrate the chamber’s wall with 3D printed clamps, enhancing the transmission of strains to the central cell culture region. This coupling enables a more uniform distribution of the traction forces through the wall and avoids the excessive deformation of the holes. The cell chamber stretching is actuated by a custom-made cell stretching system designed to be mounted on an inverted microscope stage, allowing for live cell fluorescence imaging.

### 2.2 Finite element analysis

The PDMS cell chamber was modeled as a 3D deformable solid using commercial software Abaqus 6.13.1 (SIMULIA, Dassault Systèmes). The PDMS was modeled as a neo-Hookean hyperelastic and isotropic material with Poisson’s ratio of 0.42^31^ and Young’s modulus of 2.5 MPa. The 29% strain along the y-axis of the cell culture region was achieved by the symmetrical application of a 2.65 mm displacement in the y-direction to the chamber holes. In order to accurately simulate the stretching of the cell chamber by means of the bolts, the cylindrical faces of the chamber holes were divided into two sections relative to the plane perpendicular to the stretching direction. The displacement was applied only to the sections which actually experience the bolts’ traction during stretching. Eventually, x-directional as well as y-directional logarithmic strain fields were generated (Fig. 1B). Since the cell culture region was strained by 29% uniaxially, the uniform strain field was defined as 29%±1.5% strain area, while an error of ±0.5% was considered in the transverse direction (where a negative 4.2% of strain was detected through simulations). The percentage area ratio of uniform strain fields was computed with respect to the total area of the cell culture region. Eventually, the uniform strain region was defined by superimposing the outlines of the uniform areas along x- and y-directions in Fiji software.

**Fig. 1.**
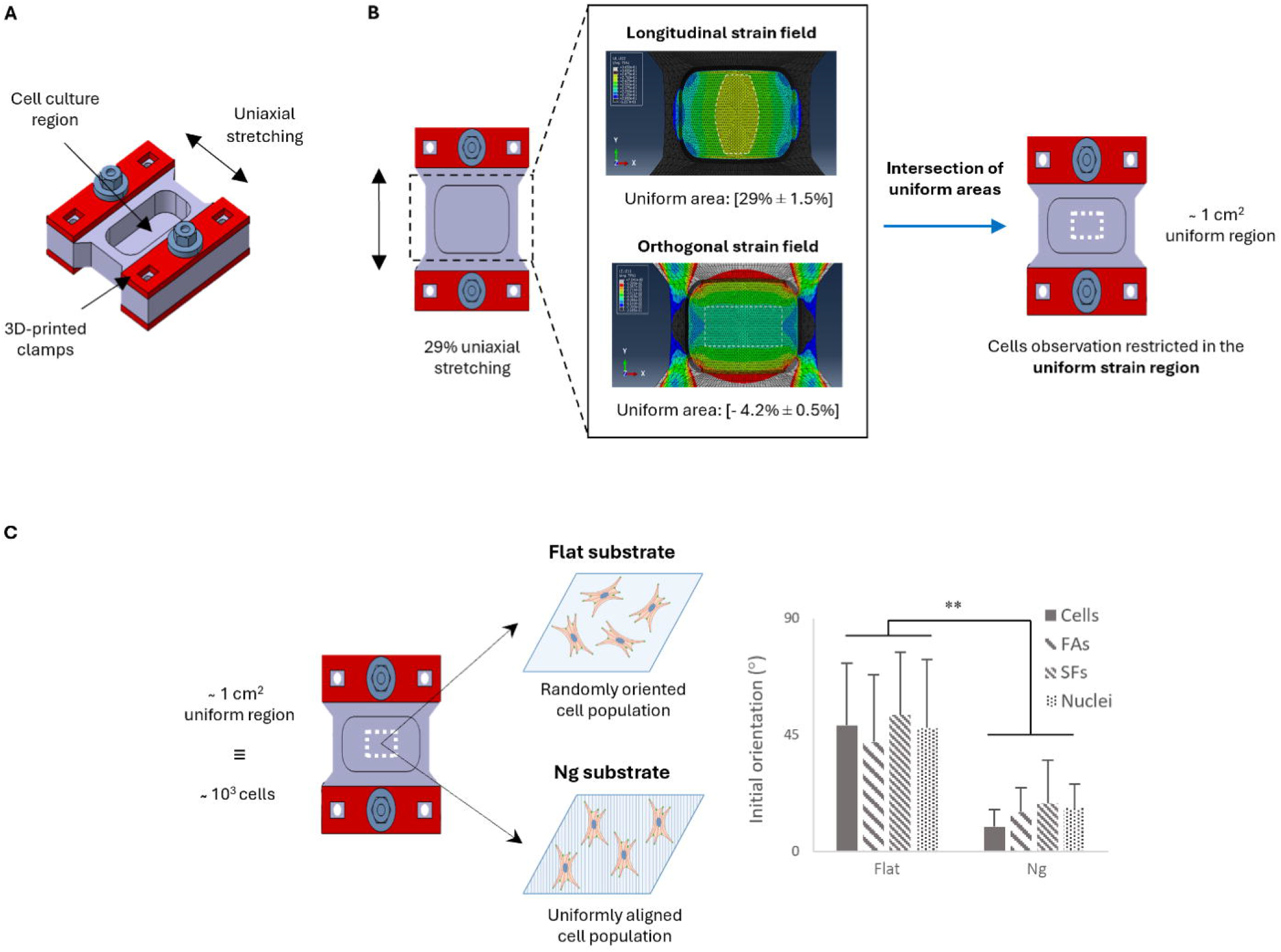
Standardization of cell orientation within a uniform uniaxial strain field. (A) Schematic of the PDMS-based cell stretching chamber mounted on a custom uniaxial device compatible with live-cell imaging. (B) Finite element analysis under 29% uniaxial stretching showing longitudinal (y) and transverse (x) logarithmic strain fields; their intersection defines the central uniform observation area. (C) Orientation distribution of NIH3T3 fibroblasts cultured for 24 h on flat and Ng PDMS substrates. Cells on flat substrates exhibit random orientation, whereas Ng substrates induce pronounced alignment along the groove direction (0°). FAs, stress fibers, and nuclei follow the same trend. Approximately 25 nuclei, 25 cells, and 140 FAs were analyzed per condition. Data are mean ± SD (*p < 0.05; **p < 0.01; n.s., not significant).

### 2.3 Cell culture

Mouse embryo fibroblasts NIH3T3 cell line were cultured at 37 °C in 5% CO2 in a humidified incubator in Dulbecco’s modified Eagle’s (DMEM) high glucose medium (Sigma-Aldrich, D5671) supplemented with 10% bovine calf serum (Sigma-Aldrich, C8056), 1% L-glutamine (Sigma, St. Louis, MO), and 1% sodium pyruvate (ThermoFisher). Prior to cell seeding, the cell stretching chamber was sterilized by ultraviolet exposure for 1 h and, then, the chamber culture surface was incubated with 50 µg/ml of human fibronectin for 1 h at 37 °C. At the end of coating procedure, cells were seeded at a density of 10^3^ cells/cm^2^ and left to adhere for 24 h in the incubator. The day after, 2 ml of cell medium were supplemented with 25 mM HEPES (ThermoFisher) and the cell chamber was mounted on the cell stretcher device.

### 2.4 Cell transfection and staining

NIH3T3 cells were seeded in a 35 mm petri dish at 80% confluency and left overnight in the incubator at 37 °C and 5% CO2. Subsequently, cells were then transiently transfected with talin-GFP or Lifeact-GFP depending on the type of experiment. The transfection complex was prepared in Opti-MEM reduced serum medium (GIBCO) and Lipofectamine 3000 (ThermoFisher) was used as a transfection reagent. The amount of DNA/Lipofectamine 3000 ratio was determined following the supplier’s instructions. Briefly, after 6 h of incubation with 2.5 μg (talin-GFP) or 1.25 μg (Lifeact-GFP) of pDNA in lipoplexes, incubation medium was replaced with complete cell culture medium. After 24 h in the incubator, cells were seeded on a cell stretching chamber and live experiments were conducted after 24 h. For cytoskeletal and nuclear analyses, cells seeded in the cell chamber were stained with Hoechst 33342 (1:10,000 in culture medium) for 20 min. Then, the incubation medium was replaced with 2 ml of complete cell culture medium supplemented with 25 mM HEPES.

### 2.5 Live cell experiments

Live cell fluorescence imaging experiments were performed on a ZEISS Axio Observer Z1. After the cell chamber was mounted on the actuation device, it was placed inside the microscope incubator where physiological environmental conditions were guaranteed. Subsequently, sustained uniaxial strains of 8% or 29% were applied to the cell culture elastic membrane. Control experiments were instead conducted in relaxed (0% strain) conditions. Cell images were acquired before, immediately (2 min) and 15 min after the application of the stretching stimulus. In particular, the 2 min time point takes into consideration the time necessary to follow cells during stretching and the time for the acquisition of the image. Fluorescence images were collected with the 63x oil immersion objective (N.A. 1.4) and preliminary experiments were carried out to optimize image settings to minimize phototoxicity and photobleaching. Live cell fluorescence experiments for cytoskeletal analysis were conducted in the same way on ZEISS LSM 700 confocal microscope. Similarly, environmental conditions inside the microscope incubator were optimized to allow live experiments and images were collected with 63x oil immersion (N.A. 1.4) objective and Z-stacks were acquired with the optimal interval suggested by the software.

### 2.6 Image analysis

Cell images analysis on flat and Ng substrates was conducted using Fiji software. Cell polarization was assessed from talin and phalloidin stained cells that were analyzed with the MomentMacroJ v1.3 script (hopkinsmedicine.org/fae/mmacro.htm) in Fiji. Briefly, Lifeact Z-stack images were Z-projected (sum intensity) and the second moment of images was calculated through the macro. For our purposes, the angle of polarization was defined as the angle that the principal axis of inertia forms with the reference axis. Nuclear images were firstly converted to binarized masks representative of their shapes using an appropriate threshold selected manually. Then, the commands “Edit”, “Selection”, “Fit Rectangle” were used to determine the minimal rectangle that encompasses the binarized images. The angle between the primary axis of rectangle and a line parallel to the x-axis of the image was defined as nuclear orientation. The major and minor axes were referred, respectively, as the major and minor edges of the fitting rectangles. The aspect ratio (AR) was then computed as the ratio of major to minor.

FA morphometric parameters were assessed following a modified procedure of the one proposed by Maruoka et al.^32^. In particular, digital images of FAs were firstly processed using “Gaussian Blur” command and then subtracted from the original images using the “Image Calculator” command. The images were further processed with “Threshold” command to obtain binarized images. The outlines of single FAs were selected with the wand tool from the digital cell images before stretching and the outlines of the same FAs were selected in the subsequent time points after stretching. The FAs morphology was extracted from the major axis and the orientation of the minimal fitting rectangle. To minimize measurement errors and obtain length variation above the image resolution, only the FAs characterized by an initial (before stretching) length (Lo) > 4 µm for control and 8% strain conditions, and Lo > 1 µm for the 29% strain condition were considered in this analysis. Stress fibers (SFs) orientation was extracted from Z-projected Lifeact confocal images through a segmentation algorithm (FSegment) implemented in Matlab. The binarized images of stress fibers were obtained, and the orientation of each stress fiber was then calculated using the “Analyze Particles” command in Fiji, by fitting the minimum enclosing ellipse. The average of all the stress fiber orientations was considered for each cell. To evaluate the fluorescence intensity of the actin cup, only a few Lifeact slides above the nucleus were Z-projected (sum intensity). On the resultant images, the mean fluorescence intensity was measured inside the outline of the nucleus.

### 2.7 Statistical analysis

All data are reported as mean ± standard deviation. Statistical comparisons were performed by means of a non-parametric Kruskal-Wallis test and P values < 0.01 (^**^) or < 0.05 (^*^) denote statistically significant differences. The Spearman Rank Correlation Coefficient (R) was implemented to assess the correlation between the datasets. Eventually, the differences in variance across the experimental conditions were evaluated through the Levene’s test.

## 3. RESULTS

### 3.1 Standardization of cell orientation with respect to controlled uniaxial stretching stimuli

Uniaxial stretching of the PDMS cell chamber was achieved using a custom-made actuation system designed to deliver controlled mechanical stimuli while enabling live-cell imaging (Fig. 1A). To calibrate the strains transmitted to the cell culture region, incremental displacements were applied to the chamber ends and the resulting longitudinal and transverse strains were quantified from fluorescent fiducial markers. The calibration curve demonstrated that the system reliably generated uniaxial strains of up to 29%, accompanied by an approximately 3.5% transverse compressive strain (Fig. S1C).

Finite element analysis (FEA) was performed to assess the spatial distribution of strain within the cell culture region under 29% uniaxial stretching (Fig. 1B). A homogeneous longitudinal strain field (29%±1,5%) was observed over 33.1% of the total culture area, while a uniform transverse strain field (-4,2%±0,5%) covered 42% of the area. Superimposition of these regions identified a central rectangular zone (12 mm x 9.3 mm, corresponding to 23.2% of the total culture area) in which both longitudinal and transverse strains were spatially uniform. Subsequent analyses of cellular response were restricted to this region to ensure consistent mechanical stimulation. To control the relative orientation between cells and the applied strain, a Ng PDMS membrane was integrated into the cell chamber such that the groove direction was aligned with the stretching axis. In particular, the Ng membrane consisted of parallel and straight channels with a groove and ridge width of 700 nm and depth of 200 nm. After 24 h of culture, NIH3T3 fibroblasts on flat substrates displayed a broad distribution of orientation angles, consistent with random orientation, whereas cells on Ng substrates exhibited a pronounced alignment along the groove direction (Fig. 1C). This alignment extended beyond the cell body to key subcellular structures, including FAs, stress fibers, and nuclei, which all exhibited preferential orientation parallel to the nanogrooves.

Therefore, Ng substrates effectively standardize cell and subcellular orientation within a spatially uniform strain field, providing a uniform reference frame for controlled and reproducible mechanical stimulation.

### 3.2 Nuclear deformation under uniaxial substrate strains

To evaluate how externally applied mechanical strains are transmitted to the nucleus, live-cell experiments were performed on NIH3T3 fibroblasts cultured on flat or Ng substrates and subjected to sustained uniaxial stretching. Nuclear morphology was analysed at two time points, 2 min and 15 min after strain application, and for two substrate strain levels (8% and 29%). Cells cultured in unstretched conditions (0% strain) served as controls (Fig. 2A). Nuclear morphology was quantified by measuring the strain of the nuclear major and minor axes (Fig. 2B-C). At 2 min after stretching, nuclei of cells aligned along the stretching direction on Ng substrates exhibited a clear elongation of the major axis and a concomitant contraction of the minor axis, with deformation magnitude increasing with substrate strain. However, nuclear deformations did not scale affinely with substrate strain: mean major axis strains of approximately 0.7% and 2.8% were observed under 8% and 29% substrate strain, respectively. In contrast, nuclei of cells cultured on flat substrates displayed a broad and heterogeneous distribution of axis deformations, but with no significant differences from control conditions when averaged across the population (Fig. 2B-D, Tab. 1). This apparent lack of response arose from the coexistence of nuclei undergoing opposite deformation modes, depending on their initial orientation relative to the stretching direction. To quantify the influence of nuclear orientation on deformation mode, we analyzed the correlation between nuclear strain metrics and initial nuclear orientation angle for both strain levels and substrate types (Fig. 2E-G). A strong negative correlation was observed between major axis strain and nuclear orientation (R = –0.83 for 8% strain, R = –0.87 for 29% strain), as well as between AR change and nuclear orientation (R = –0.77 for 8% strain, R = –0.73 for 29% strain). Conversely, minor axis strain showed a positive correlation with nuclear orientation (R = 0.71 for 8% strain, R = 0.84 for 29% strain). Thus, nuclei aligned parallel to the stretching direction elongated along their major axis, whereas nuclei oriented perpendicular to the strain direction underwent contraction along the same axis. For clarity, cells on flat substrates were subdivided into three orientation sectors containing equal numbers of nuclei, and nuclear deformation parameters were analysed for each group under 29% substrate strain (Fig. 2H-L). Nuclei oriented parallel to the stretching direction exhibited the largest major-axis elongation, the greatest minor-axis contraction, and the highest increase in AR. In contrast, nuclei oriented perpendicularly to the stretching direction showed opposite deformations, resulting in a minimal AR change. Intermediate orientations instead displayed intermediate deformation levels. At the population level, this orientation dependence led to a marked difference between flat and Ng substrates. On flat substrates, the random distribution of nuclear orientation resulted in deformation modes that largely cancel out when averaged, yielding near-zero strain values. On Ng substrates, where nuclei were uniformly aligned with the stretching direction, nuclear deformation was coherent and directionally consistent across the population.

**Fig. 2.**
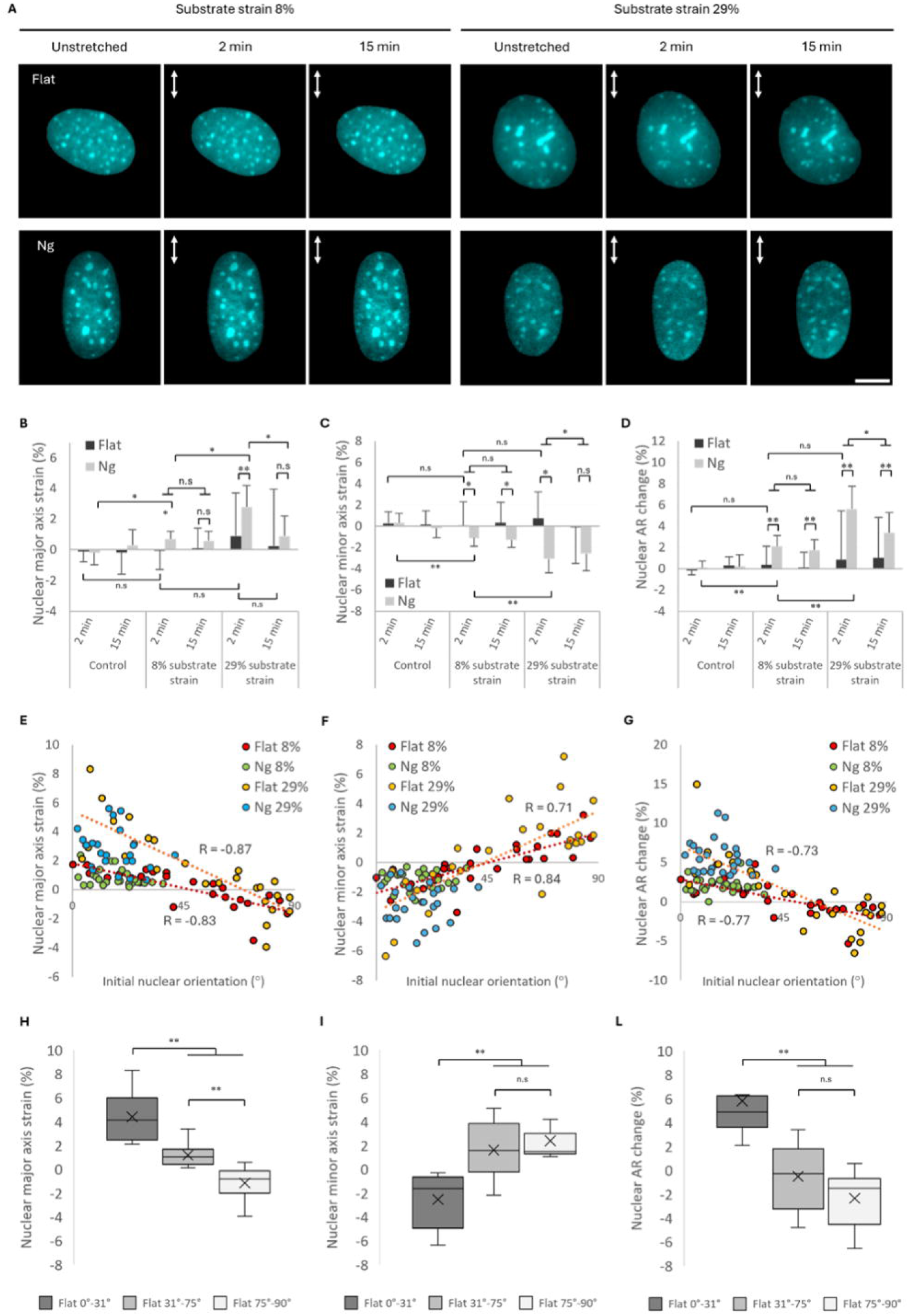
Nuclear deformation under uniaxial substrate strains. (A) Representative fluorescence images of NIH3T3 nuclei on flat and Ng substrates under unstretched conditions and after application of 8% and 29% uniaxial strain at 2 min and 15 min. White arrows indicate the stretching direction (scale bar, 15 µm). (B–C) Quantification of nuclear major axis strain (B) and minor axis strain (C) as a function of substrate strain, substrate type, and time. (D) Changes in nuclear aspect ratio (AR) following stretching. (E–G) Correlation between initial nuclear orientation and major axis strain (E), AR change (F), and minor axis strain (G) at 2 min. (H–L) Nuclear deformation parameters for flat-substrate cells subdivided into three orientation sectors under 29% strain. Approximately 25 nuclei were analyzed per condition across five independent experiments; controls included 15 nuclei per substrate. Data are mean ± SD (*p < 0.05; **p < 0.01; n.s., not significant).

Analysis at 15 min after strain application revealed a strain-dependent temporal response. Under 8% sustained strain, nuclear deformation remained comparable to that observed at 2 min for both flat and Ng substrates. In contrast, cells subjected to 29% strain exhibited a significant reduction in nuclear major-axis elongation and a corresponding increase in the minor-axis length over time (Fig. 2B-C), indicating a partial relaxation of nuclear deformation.

Taken together, these results highlight that nuclear deformation under uniaxial stretching is strongly orientation dependent and becomes uniform and quantifiable only when cell and nuclear alignment is controlled by substrate topography.

### 3.3 Deformation at the cellular level under uniaxial substrate strains

To determine how substrate deformation is transmitted to adherent cells, the strain of cellular major and minor axes was quantified following uniaxial stretching of flat and Ng substrates (Fig. 3A-B). At 2 min after strain application, cells cultured on Ng substrates exhibited affine deformation with respect to the applied substrate strain, characterized by elongation of the major axis and reduction of the minor axis at both 8% and 29% strain levels. In contrast, cells cultured on flat substrates displayed significantly attenuated deformation, with mean axis strains that were markedly lower than those observed on Ng substrates (p < 0.01). When cellular axis strains were plotted as a function of initial cell orientation, a strong inverse correlation was observed between orientation angle and major-axis strain, and a corresponding positive correlation with minor-axis strain, for both substrate strain levels (Fig. 3D–E). Changes in cellular aspect ratio (AR) were used as an integrated measure of cell deformation (Fig. 3C). Cells on Ng substrates exhibited a significant increase in AR relative to unstretched controls (9.8% and 33.9% for 8% and 29% strain, respectively), whereas no significant AR change was detected for cells on flat substrates. Orientation-resolved analysis revealed that AR changes decreased progressively with increasing misalignment between the cell major axis and the stretching direction (Fig. 3F). To assess variability in the cellular response, Levene’s test was applied to compare variance across experimental conditions (Tab. 1). Cells on flat substrates exhibited significantly higher variance in axis strain and AR change compared to those on Ng substrates (p < 0.01), indicating greater heterogeneity in deformation. Subdivision of flat-substrate cells into orientation sectors under 29% strain further confirmed that cells aligned parallel to the stretching direction experienced maximal deformation, whereas perpendicularly oriented cells exhibited minimal net AR change (Fig. 3G–I). At 15 min after strain application, cells on Ng substrates subjected to 29% strain showed a significant reduction in both major and minor axis strain compared to the 2 min time point (p < 0.05), whereas cells exposed to 8% strain maintained their initial deformation (Fig. 3A– B).

**Fig. 3.**
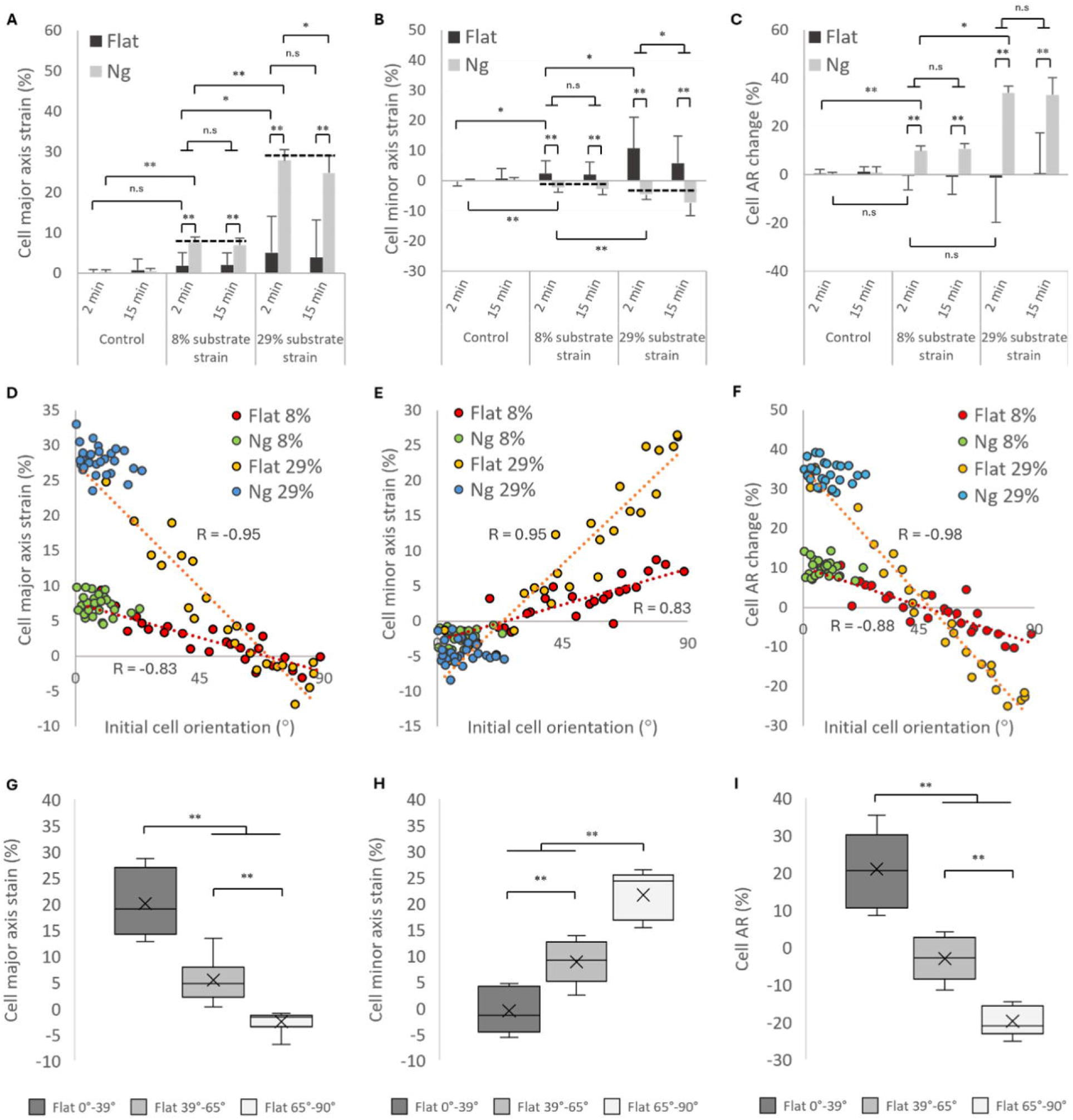
Transmission of uniaxial substrate strains at the cellular level. (A–B) Quantification of cell major-axis strain (A) and minor-axis strain (B) following 8% and 29% uniaxial substrate strain on flat and Ng substrates at 2 min and 15 min. Black dotted lines indicate substrate strains estimated from membrane calibration. (C) Changes in cellular aspect ratio (AR) as an integrated measure of deformation. (D–E) Correlation between initial cell orientation and major-axis strain (D) and minor-axis strain (E) at 2 min. (F) AR change as a function of initial cell orientation. (G–I) Deformation parameters for flat-substrate cells subdivided into three orientation sectors under 29% strain. Approximately 25 cells were analyzed per condition across seven independent experiments; controls included 10 cells per substrate. Data are mean ± SD (*p < 0.05; **p < 0.01; n.s., not significant).

These results show that substrate-to-cell strain transmission is strongly orientation dependent and becomes uniform and affine only when cell alignment is imposed by substrate nanotopography.

### 3.4 FA dynamic response under uniaxial substrate strains

The FA response to mechanical stimulation was examined in NIH3T3 cells expressing Talin-GFP and cultured on flat or Ng substrates (Fig. 4A). FA length strain was quantified by tracking individual adhesions before and after uniaxial stretching. At 2 min after strain application, FA length increased significantly as a function of substrate strain for both culture conditions (p < 0.01; Fig. 4B). On Ng substrates, FA length strain closely matched the applied substrate strain, reaching mean values of 7.2% and 28.3% under 8% and 29% substrate strain, respectively. In contrast, FAs on flat substrates exhibited significantly lower mean length strains (3.1% and 16.4%, respectively; p < 0.01). Orientation-resolved analysis revealed a strong inverse correlation between FA length strain and initial FA orientation angle (Fig. 4C). FAs aligned parallel to the stretching direction experienced maximal elongation, whereas those oriented perpendicular to the strain direction exhibited minimal deformation. This orientation dependence resulted in significantly greater variability in FA strain on flat substrates compared to Ng substrates, as confirmed by variance analysis (Tab. 1). Subdivision of FAs on flat substrates into angular sectors under 29% strain further illustrated this heterogeneity: adhesions aligned with the stretching direction exhibited the greatest elongation, intermediate orientations displayed moderate strain, and perpendicularly oriented FAs showed minimal response (Fig. 4D). At 15 min after strain application, FA length strain remained stable under 8% substrate strain for both substrate types. In contrast, FA length decreased significantly under 29% strain (p < 0.05), indicating partial relaxation of FA deformation over time. Overall, these data demonstrate that FA deformation under uniaxial stretching is affine and uniform only when adhesion orientation is constrained by Ng substrates.

**Fig. 4.**
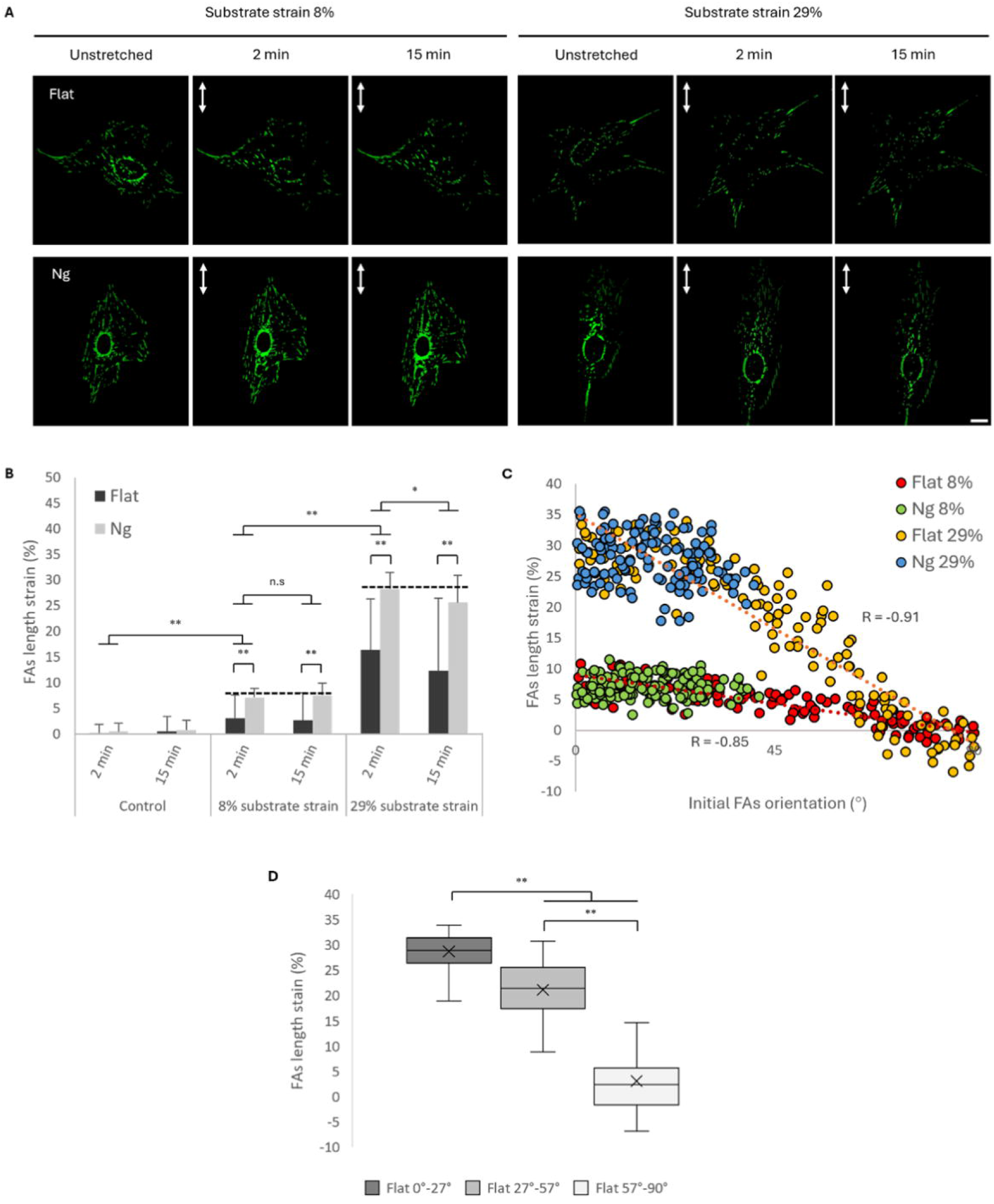
FA deformation under uniaxial substrate strain. (A) Representative fluorescence images of focal adhesions (Talin-GFP) in NIH3T3 cells cultured on flat and Ng substrates before and after uniaxial stretching (scale bar, 20 µm). (B) Average FA length strain at 2 min and 15 min following application of 8% and 29% substrate strain. (C) FA length strain as a function of initial FA orientation relative to the stretching direction at 2 min. (D) FA length strain for flat-substrate cells subdivided into three orientation sectors under 29% strain. FAs aligned with the stretching direction (0°–27°) exhibited maximal elongation, whereas those nearly perpendicular (57°–90°) showed minimal deformation; intermediate orientations (27°–57°) displayed intermediate strain. Approximately 140 FAs were analyzed per condition across 10 cells and five independent experiments; controls included 50 FAs per substrate. Data are mean ± SD (*p < 0.05; **p < 0.01; n.s., not significant).

### 3.5 Cytoskeletal adaptation under uniaxial substrate strains

To assess how externally applied forces, perceived through FAs, are integrated and distributed within the cell, quantitative analyses were conducted on live cells transfected with Lifeact and stained for the nucleus with Hoechst 33342 (Fig. 5A). Immediately after strain application, stress fibers reoriented towards the stretching direction, with the magnitude of reorientation increasing with substrate strain level (Fig. 5B). Cells cultured on flat substrates showed significantly larger angular reorientation compared to those cultured on Ng substrates (p < 0.01), reflecting the initially random orientation in the absence of topographical guidance. These orientation changes persisted over time, with no significant differences between 2 min and 15 min time points. To assess strain-dependent redistribution of cytoskeleton tension, changes in actin-cap fluorescence intensity were quantified (Fig. 5C). Actin-cap intensity increased significantly with substrate strain magnitude (p < 0.01), and decreased over time (p < 0.05), consistent with dynamic remodelling of cytoskeletal structure under sustained loading. No significant differences in actin-cap intensity changes were detected between flat and Ng substrates.

**Fig. 5.**
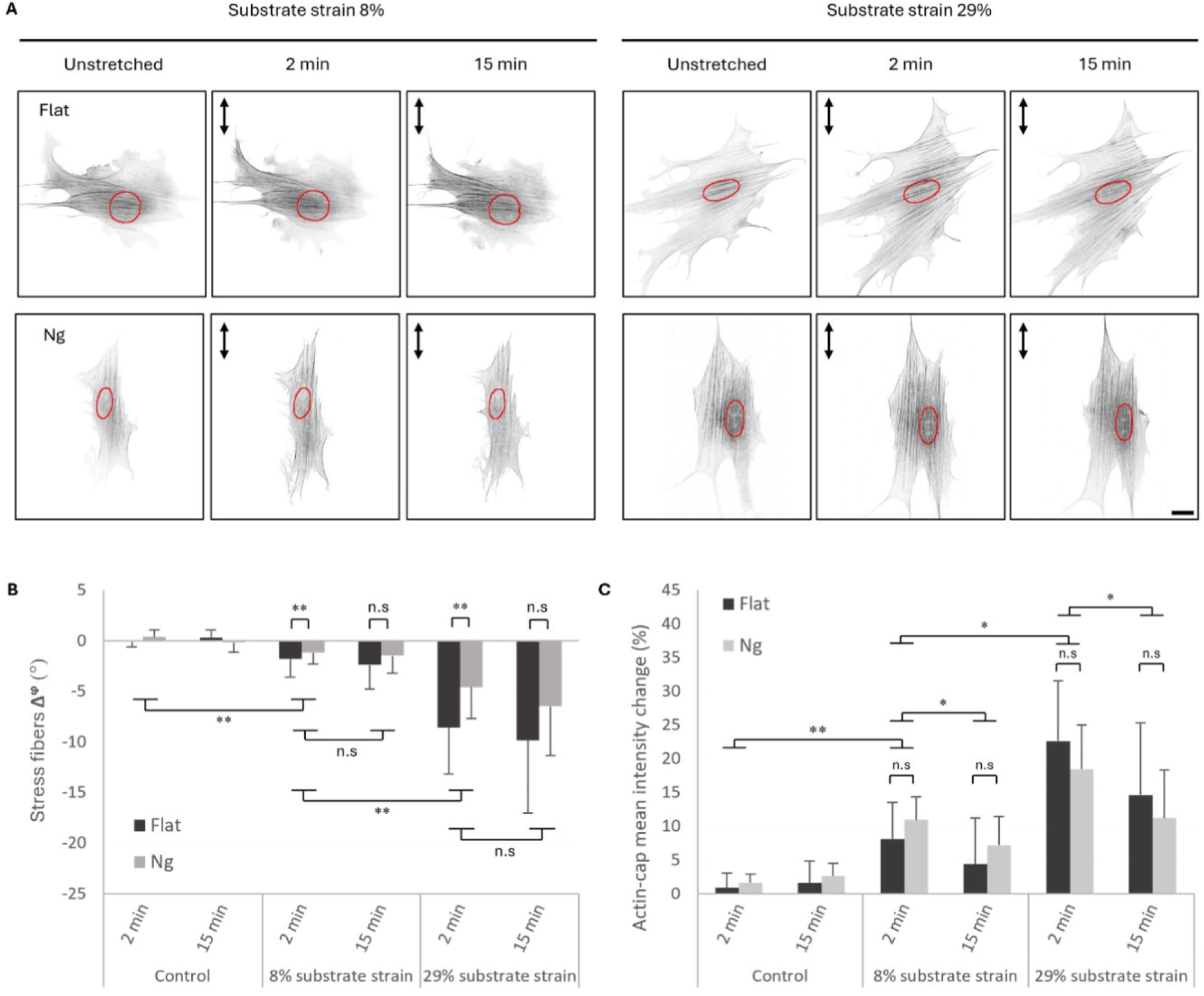
Cytoskeletal response under uniaxial substrate strains. (A) Representative confocal fluorescence images of the actin cytoskeleton (Lifeact-GFP) and nuclei (red) in NIH3T3 cells cultured on flat and Ng substrates before and after uniaxial stretching (scale bar, 10 µm). (B) Average change in stress fiber orientation (Δφ) following application of 8% and 29% substrate strain at 2 min and 15 min. (C) Changes in actin-cap mean fluorescence intensity as a function of substrate strain and time. Approximately five cells were analyzed per condition across three independent experiments. Data are mean ± SD (*p < 0.05; **p < 0.01; n.s., not significant).

These results indicate that cytoskeletal reorganization and actin-cap recruitment are primarily governed by strain magnitude rather than substrate-induced alignment.

## 5. Discussion

Mechanical stimulation is increasingly recognized as a fundamental regulator of cellular behavior in both physiological and pathological contexts^33^. Accordingly, a wide range of devices and platforms have been developed to apply controlled mechanical forces to cells in vitro, with the aim of directing processes such as differentiation, migration, and proliferation. While substantial effort has been devoted to refining the precision and tunability of these systems, comparatively less attention has been paid to how uniformly individual cells within a population perceive the applied mechanical stimulus^17^.

In this study, we demonstrated that the consistency of mechanical signal perception at the single-cell level represents a critical, yet unappreciated, determinant of mechanobiological outcome. Specifically, we showed that the relative orientation between cells and the direction of applied strain introduces a major source of variability in force transmission, which fundamentally limits the reproducibility of population-level measurements when flat substrates are used. Using this framework, we systematically dissected how substrate-applied strains are propagated across multiple cellular scales. At the level of FAs, nanogroove-confined adhesions exhibited substrate affine elongation immediately after stretching, closely matching the magnitude of applied strain. To our knowledge, such direct correspondence between substrate deformation and FA length change has not been previously reported. This result highlights the importance of controlling adhesion orientation when quantifying force transmission in mechanobiology experiments. At the cellular scale, aligned cells on Ng substrates underwent affine deformation in response to axial stretching, whereas cells on flat substrates displayed significantly attenuated and highly heterogeneous responses. Orientation-dependent analyses revealed that cells aligned parallel to the stretching direction experienced maximal elongation, while those oriented perpendicularly underwent relative compression. On flat substrates, the random distribution of cell orientations resulted in population-averaged responses that were close to zero, despite pronounced deformation occurring at the single-cell level. This orientation-induced cancellation effect was particularly evident at the nuclear level. Nuclear deformation emerged as a highly sensitive and functionally relevant readout of mechanical stimulation, yet only when cell orientation was controlled. On Ng substrates, nuclei consistently elongated along their major axis and reduced their minor axis in response to stretching, resulting in coherent increases in nuclear AR across the population. In contrast, nuclei on flat substrates exhibited opposing deformation modes depending on their initial orientation, leading to negligible average changes and increased variability. The observation that nuclear deformation did not scale affinely with substrate strain, even under aligned conditions, highlights the intrinsic mechanical resistance of the nucleus. Previous studies have shown that the nucleus is substantially stiffer than the surrounding cytoplasm^34^ and that its deformability is modulated by prestress generated by the cytoskeleton^35^. In this context, the actin cap plays a pivotal role by transmitting cytoskeletal forces directly to the nuclear envelope^36^. Consistent with this mechanism, we observed strain-dependent recruitment of actin to the actin cap, indicating increased cytoskeletal tension acting on the nucleus following substrate stretching. Notably, actin-cap reinforcement occurred to a similar extent on both flat and Ng substrates, yet significant nuclear deformation was observed only in aligned cells. This finding emphasizes that the magnitude of intracellular force alone is insufficient to predict nuclear deformation; rather, the relative orientation between force-generating cytoskeletal structures and the nucleus is a decisive factor. These results agree with previous reports showing that nuclear deformation under substrate stretching is strongly orientation dependent and is abolished when stress fiber tension is pharmacologically reduced^26^.

Beyond the immediate mechanical response, our results revealed also a strain-dependent adaptive behavior of cells subjected to sustained uniaxial stretching. Moderate strains (8%) induced persistent morphometric changes at the focal adhesion, cytoskeletal, and nuclear levels, indicative of stable mechano-adaptation. Such responses are consistent with previous studies reporting enhanced actin polymerization, focal adhesion maturation, and reinforcement of cell–substrate coupling under physiological strain levels^37–39^. In contrast, exposure to high strain levels (29%) triggered a transient response characterized by partial relaxation of cellular and subcellular deformations within 15 min. This behavior is consistent with the activation of protective mechanisms aimed at limiting excessive intracellular tension and preserving structural integrity, such as cytoskeletal remodelling and force shedding^40–43^. Importantly, this strain-threshold–dependent adaptation was observed on both flat and nanogrooved substrates, indicating that while alignment governs the uniformity of force transmission, the intrinsic adaptive response is primarily dictated by strain magnitude.

Collectively, these findings highlight that substrate nanotopography does more than simply align cells: it fundamentally reduces cell-to-cell variability in mechanical signal perception. On flat substrates, uncontrolled orientation leads to divergent mechanosensory activation, heterogeneous focal adhesion dynamics, and incoherent nuclear deformation, thereby obscuring meaningful mechanobiological relationships. By contrast, nanogroove-guided alignment enables population-wide coherence in mechanical response, allowing robust interpretation of how specific strain magnitudes modulate cellular mechanics.

From a broader perspective, the ability to standardize mechanical dose delivery has important implications for both basic mechanobiology and applied biomaterials research. Reducing variability in mechanical stimulation is essential for elucidating causal links between physical cues and downstream biological outcomes, including gene regulation and cell fate decisions. Moreover, integrating topographical guidance with mechanical loading provides a versatile design principle for engineered tissues that must withstand and respond to dynamic mechanical environments, such as cardiac muscle, tendons, and vascular tissues^44–47^.

## 6. Conclusion

In this study, we demonstrated that controlling cell orientation is a critical prerequisite for achieving consistent and reproducible mechanical stimulation in vitro. By integrating Ng substrate topography with uniaxial stretching, we established a simple yet effective strategy to standardize the mechanical dose delivered to cells across an entire population. Nanogroove-guided alignment enabled the coherent transmission of externally applied strains from the substrate to the focal adhesions, the cytoskeleton, and ultimately the nucleus. As a result, nuclear deformation, that is probably the most functionally relevant readout of mechanotransduction, became uniform and quantifiable at the population level, in contrast to the heterogeneous and orientation-dependent responses observed on conventional flat substrates.

Our findings further revealed that cellular adaptation to sustained mechanical loading is governed primarily by strain magnitude rather than substrate topography, highlighting the existence of effective mechanical stimulation windows that promote stable mechano-adaptive responses. Importantly, while nanotopography does not alter intrinsic cellular adaptation mechanisms, it eliminates cell-to-cell variability in mechanical signal perception, thereby uncovering mechanobiological responses that are otherwise masked in standard culture systems.

This work establishes substrate-guided cell alignment as a robust and broadly applicable strategy to standardize mechanical dose delivery in mechanobiology experiments. By reducing response heterogeneity and enabling reproducible nuclear mechanostimulation, this approach provides a practical framework for improving experimental reliability and for advancing the development of mechanomedicine, tissue engineering, and biomaterial-based mechanotransductive platforms.

**Table 1.** Variance analysis of cellular and subcellular deformation parameters. Results of Levene’s test assessing differences in variance across experimental conditions for cellular and subcellular deformation parameters, including cell axes strain, FA length strain, stress fiber reorientation, and nuclear deformation metrics. Statistical significance indicates increased heterogeneity in response between flat and Ng substrates.

|  | Control |  |  |  | 8% substrate strain |  |  |  | 29% substrate strain |  |  |  |
| --- | --- | --- | --- | --- | --- | --- | --- | --- | --- | --- | --- | --- |
|  | 2 min |  | 15 min |  | 2 min |  | 15 min |  | 2 min |  | 15 min |  |
|  | Flat | Ng | Flat | Ng | Flat | Ng | Flat | Ng | Flat | Ng | Flat | Ng |
| Cell major axis strain (%) | n.s |  | n.s |  | p < 0.01 |  | n.s |  | p < 0.01 |  | n.s |  |
| Cell minor axis strain (%) | n.s |  | n.s |  | p < 0.01 |  | p < 0.01 |  | p < 0.01 |  | p < 0.01 |  |
| Cell AR change (%) | n.s |  | n.s |  | p < 0.01 |  | p < 0.01 |  | p < 0.01 |  | p < 0.01 |  |
| FAs length strain (%) | n.s |  | n.s |  | p < 0.01 |  | p < 0.01 |  | p < 0.01 |  | p < 0.01 |  |
| Stress fibers $\Delta f$ | n.s | | n.s | | n.s | | n.s | | n.s | | n.s | |
| Actin-cap mean intensity change (%) | n.s |  | n.s |  | n.s |  | n.s |  | n.s |  | n.s |  |
| Nuclear major axis strain (%) | n.s |  | n.s |  | p < 0.05 |  | p < 0.01 |  | p < 0.05 |  | p < 0.01 |  |
| Nuclear minor axis strain (%) | n.s |  | n.s |  | p < 0.01 |  | p < 0.01 |  | p < 0.01 |  | p < 0.05 |  |
| Nuclear AR change (%) | n.s |  | n.s |  | p < 0.01 |  | p < 0.05 |  | p < 0.01 |  | p < 0.05 |  |

