## Supplementary material for "Standardizing mechanical dose delivery to cells via nanogroove-guided alignment": S1

**Experimental design**

Cell response to uniaxial substrate stretching on flat and nanogrooved (Ng) substrates was investigated by live-cell fluorescence imaging to resolve both the immediate mechanical response and the short-term adaptive behavior of cells and their mechanosensory components (focal adhesions (FAs), cytoskeleton, and nucleus). Imaging was performed at two time points: 2 min and 15 min after strain application. The 2 min time point, dictated by the technical time required for image acquision, was selected to capture the immediate response of cells following substrate deformation. The 15 min time point was chosen to probe the short-term cellular adaptation to sustained strain, within a time window compatible with FA assembly–disassembly dynamics and cytoskeletal remodeling[1,2]. Sustained uniaxial strains of 8% and 29% were applied using a custom-built stretching system. The first strain magnitude was selected based on literature evidence indicating a lower threshold of 3% for cellular mechanosensing and mechanoresponsiveness[3], as well as on the physiological strain range (5–15%) experienced by most tissues[4-6]. Conversely, the 29% strain level was chosen to account for the large deformations (>20%) undergone by specific tissues, such as the solid tissues of hollow organs (e.g., bladder and stomach) and the skin[7,8]. Moreover, although significant pathology can arise if relatively large stretches are sustained[9,10], the behaviour of cells is still not known at these levels of sustained strain.

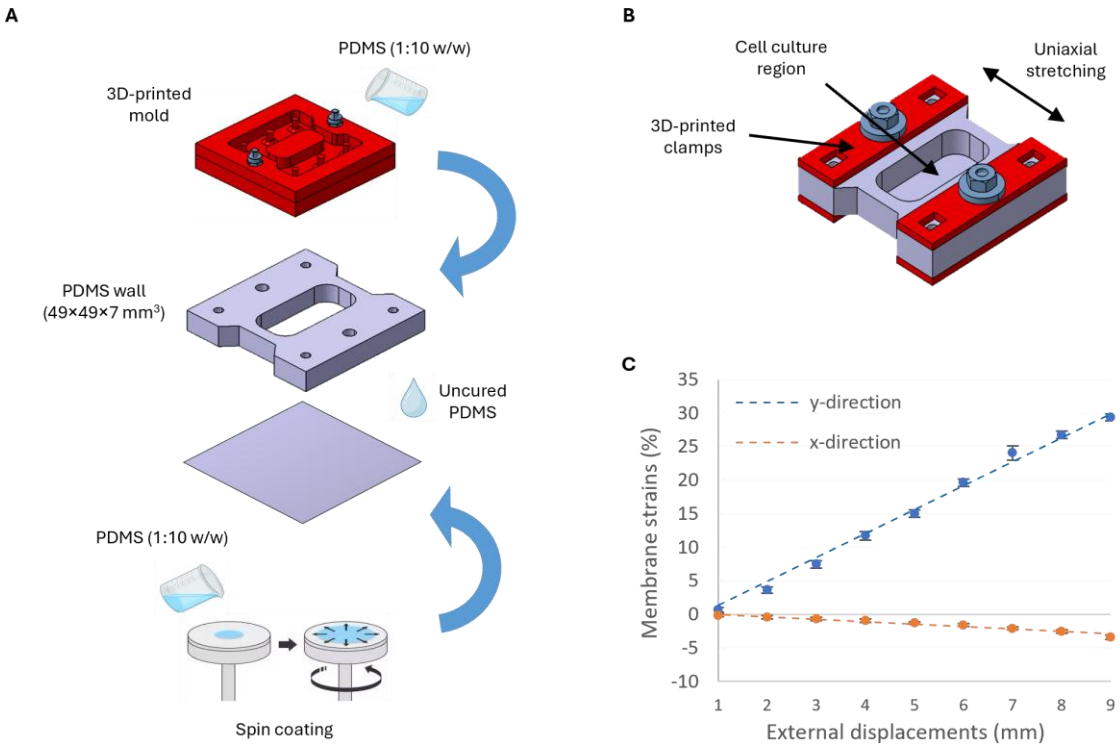


**Figure S1.** Fabrication and calibration of cell stretching chamber (A) Fabrication workflow of the cell chamber. The chamber was entirely made of polydimethylsiloxane (PDMS), prepared by mixing base and curing agent at a 10:1 weight ratio. Flat and Ng membranes with a thickness of ~100 µm were obtained by spin-coating the PDMS mixture at 500 rpm for 1 min. The chamber well was fabricated by casting PDMS in a 3D-printed master and curing at 80 °C for 2 h. The membrane was bonded to the chamber well using uncured PDMS as an adhesive layer, followed by a second curing step at 80 °C for 2 h to obtain the assembled device. (B) Schematic of the optimized PDMS cell-stretching chamber for uniaxial deformation. The device (49 × 49 × 7 mm³) features a central bottom region (32 × 15 × 0.1 mm³) where cells are cultured. Uniaxial stretching is applied symmetrically by pulling the chamber at its two ends. Each end contains three holes: the two lateral holes are used for traction, while the central hole is coupled to 3D-printed clamps that integrate with the chamber walls, improving strain transmission to the cell-culture area. This design ensures a more uniform force distribution and prevents excessive deformation of the holes. (C) Calibration of the uniaxial strains transmitted to the cell culture region. Longitudinal and transverse strains were calculated by tracking fluorescent markers on the membrane in response to incremental uniaxial displacements of 1 mm applied to the chamber holes via the actuation system. The calibration of membrane strains was performed on 2 different cell chambers (n=2).

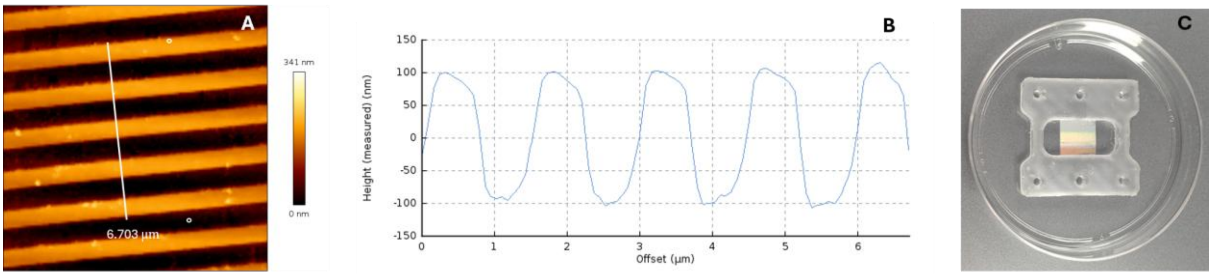


**Figure S2.** Characterization of Ng membranes. (A) Representative atomic force microscopy (AFM) topography image of the PDMS replica obtained from the nanopatterned master, showing well-defined and regularly spaced nanogrooves. The pattern consisted of parallel nanogrooves with a groove and ridge width of 700 nm and a depth of 200 nm. (B) Height profile extracted along the white line indicated in (A), confirming faithful replication of the nanogroove geometry. AFM measurements were performed using a NanoWizard Pure system operated in Quantitative Imaging (QI) mode with FESP-V2 silicon cantilevers. The setpoint was 0.5 V, the Z-length was 300 nm, and the Z-speed was 150 µm/s. Images were acquired with a lateral resolution of 0.039 µm per pixel. (C) Final PDMS cell-stretching chamber with the Ng membrane bonded to the chamber wall, forming the complete device for cell culture and uniaxial mechanical stimulation.

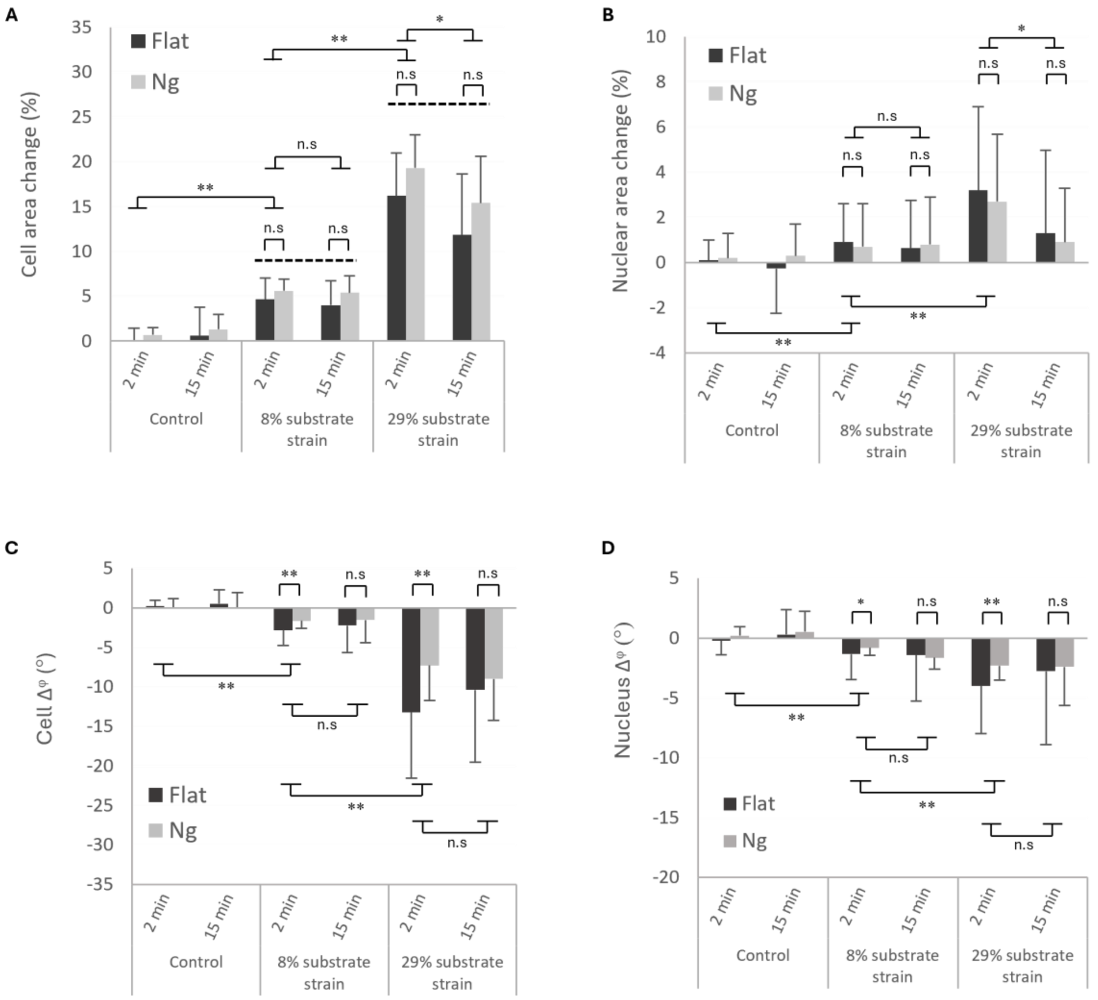


**Figure S3.** Cellular and nuclear area changes and reorientation under uniaxial substrate strains. Changes in cell and nuclear area and orientation were quantified for NIH3T3 cells cultured on flat and Ng substrates following the application of 8% and 29% uniaxial strain, evaluated 2 min and 15 min after stretching. For orientation analyses, the nanopattern direction was defined as 0° and angles were reported within [0°, 90°]. Orientation shifts ($\Delta^{\varphi}$) were calculated for each cell as the difference between the orientation after stretching at each time point and the initial orientation. (A) Percentage changes in cell spreading area compared with the expected substrate area changes derived from the calibration (dotted lines, 7.1% and 25.1% for 8% and 29% strain, respectively). Cell spreading area increased immediately after stretching in a strain-dependent manner ($p < 0.01$) with no significant differences between the two substrate types. However, this increase was lower than that expected from substrate deformation, consistent with the discrete nature of cell-substrate interaction which is mediated by focal contacts. After 15 min of 8% sustained strain, cell area returned to control values, whereas a significant reduction over time was observed at 29% strain ($p < 0.05$). (B) Average change in cell body orientation ($\Delta^{\varphi}$) following stretching. Randomly oriented cells on flat substrates underwent a significantly larger reorientation toward the strain direction than cells pre-aligned on Ng substrates ($p < 0.01$). (C) Changes in nuclear spreading area. No significant differences were observed between flat and Ng substrates. Nuclear area increased proportionally to strain level ($p < 0.01$) and slightly decreased after 15 min under 29% strain ($p < 0.05$). (D) Average change in nuclear orientation ($\Delta^{\varphi}$). Nuclei on flat substrates showed a significantly larger reorientation in the direction of strain than those on Ng substrates. In both conditions, nuclear reorientation increased with strain level and remained approximately constant over time. For each strain level and substrate type, approximately n = 25 cells and n = 25 nuclei were analyzed over time. Data are reported as mean ± SD. Statistical significance (Kruskal-Wallis test) is indicated (*$p < 0.05$; **$p < 0.01$; n.s., not significant).
